# Adhesion to type I collagen fibrous gels induces E- to N-cadherin switching without other EMT-related phenotypes in lung carcinoma cell A549

**DOI:** 10.1101/2020.10.02.323881

**Authors:** Hitomi Fujisaki, Sugiko Futaki, Masashi Yamada, Kiyotoshi Sekiguchi, Toshihiko Hayashi, Shunji Hattori

## Abstract

In culture system, environmental factors, such as increasing exogenous growth factors and adhesion to type I collagen (Col-I) induce epithelial-to-mesenchymal transition (EMT) in cells. Col-I molecules maintain a non-fibril form under acidic conditions, and they reassemble into fibrils under physiological conditions. Col-I fibrils often assemble to form three-dimensional gels. The gels and non-gel-form of Col-I can be utilized as culture substrates and different gel-forming state often elicit different cell behaviors. However, gel-form dependent effects on cell behaviors, including EMT induction, remain unclear. EMT induction in lung cancer cell line A549 has been reported via adhesion to Col-I but the effects of gel form dependency are unelucidated. This study investigated the changes in EMT-related behaviors in A549 cells cultured on Col-I gels.

We examined cell morphology, proliferation, single-cell migration and expression of EMT-related features in A549 cells cultured on gels or non-gel form of Col-I and non-treated dish with or without transforming growth factor (TGF)-β1. On Col-I gels, some cells kept cell–cell contacts and formed clusters, others maintained single-cell form. In cell–cell contact regions, E-cadherin expression was downregulated, whereas that of N-cadherin was upregulated. Vimentin and integrins α2 and β1 expression were not increased. In TGF-β1-treated A549 cells, cadherin switched from E- to N-cadherin. Their morphology changed to a mesenchymal form and cells scattered with no cluster formation. Vimentin, integrins α2 and β1 expression were upregulated. Thus, we concluded that culture on Col-I fibrous gels induced E- to N-cadherin switching without other EMT-related phenotypes in A549 cells.

## Introduction

Type I collagen (Col-I) is a ubiquitous and major component of interstitial tissues. *In vivo*, many kinds of cells are found resting proliferation within Col-I fibrils containing matrix, while, cell– extracellular matrix (ECM) interactions are essential for cell movement (1). For a long while, Col-I has been widely studied and used as a cell culture substrate (1–7). Typically, Col-I is solubilized under acidic conditions and extracted from animal tissues. Col-I remains in a non-fibril form under acidic conditions, meaning that Col-I molecules are not assembled each other. Soluble Col-I molecules reassemble into fibrils under physiological conditions (i.e., pH, temperature, and ionic strength) spontaneously, and these fibrils often assemble into fiber bundles with a three-dimensional structure (i.e., gels) (2–4). Col-I, as a culture equipment, has been roughly divided into two categories depending on whether or not gels are formed. In 1982, Kleinman et al. reported a few different Col-I treatment protocols for culture substrates, such as drying up acidic Col-I solution to cover the surface of a culture dish (or removing excess solution) and neutralizing acidic Col-I solution for gelation (2, 3). They called the former substrate “collagen films” and the latter “reconstituted collagen” (3). Sometimes, the former is called “2D culture substrate” and the latter is “3D culture substrate” respectively. Recently fibril-forming Col-I has been used as novel hydrogels (5) and the history of the collagen gel culture assay has been reviewed (6).

Adherence to different forms of Col-I (gels or non-gel-form) elicits different cellular responses in various cells, such as morphological appearance and metabolic and proliferative potential. In fibroblast culture, there are differences between gels (3, 7) and non-gel-form (3) in terms of the growth rates, mitogenic responses to growth factors, and collagen synthesis. Koyama *et al*. showed that arterial smooth muscle cells remained arrested in G1 phase in the presence of Col-I gels. However, when cultured on non-gel-form of Col-I, they proliferate (8). Furthermore, the culture of human keratinocytes on the non-gel form of Col-I results in the spread morphology and continuous growth. While, the culture on Col-I gels induces round morphology and apoptosis in cells (9). The level of reactive oxygen species (ROS) in 3T3-L1 murine preadipocytes increases on Col-I, independent of the gel-forming state, compared with on a non-treated dish. A markedly higher level of ROS is found in cells on Col-I gels, it serves as a suppressor of cell proliferation and migration. But higher level of ROS acts as a positive regulator in these processes on the non-gel-form of Col-I. These opposite effects of increasing ROS are attributed to the levels of nuclear factor-κB/p65 activation (10).

Epithelial-to-mesenchymal transition (EMT) is a fundamental biological process, whereby epithelial cells lose their apical-basal polarity, cell–cell adhesions via E-cadherin weaken, and the cells express mesenchymal cell characteristics, including N-cadherin expression (11–13). Beside the switching of cadherin types, various EMT-related markers (i.e., vimentin, matrix metalloproteinases, and transcription factors Snail, Slug, and Twist1) have been reported (12–15). The concept of EMT was proposed by Greenburg and Hay as a process induced by adherence to Col-I gels in embryonic chick corneal or lens epithelial cells (16). More recently, EMT has been associated with wound healing and cancer metastasis (17–20). Since the turn of the century, the number of papers on EMT has increased exponentially, and there are also more reports about intermediate cell behaviors between the epithelial and mesenchymal forms. The EMT International Association proposed naming the intermediate form, epithelial mesenchymal plasticity (EMP) (12). Many reports and discussions are ongoing regarding EMT and EMP. Various growth and differentiation factors can induce and regulate the EMT process. Among them, transforming growth factor (TGF)-β is one of the well-characterized inducers or enhancers of EMT (21–24). Numerous signaling pathways that regulate or mediate the EMT process via the activation of TGF-β receptors have been reported. Furthermore, increased exogenous TGF-β upregulates ECM production and deposition, including Col-I, around the cellular microenvironment (23, 24). Col-I deposition has been attributed to increasing matrix stiffness. Additionally, TGF-β1 signal activation contributes to a positive feedback autocrine loop in tumor cells undergoing EMT (25).

Adhesion to Col-I induces EMT-like upregulation of single-cell motility by regulating cellular signal transduction (26–29). Tiam1/Rac signaling promotes motility in Madin–Darby canine kidney cells cultured on Col-I, which is regulated by phosphatidylinositol 3-kinase (PI3K) pathway activation (26). Cells converted to highly invasive forms of pancreatic cancer are characterized by extensive Col-I deposition (27, 28). Shintani *et al*. showed that mouse mammary epithelial cells upregulated N-cadherin expression and underwent EMT in response to culture dishes coated with Col-I through a pathway involving integrins and discoidin domain receptor tyrosine kinase 1, including the activation of PI3K, Rac1, and c-Jun N-terminal kinases. These effects are not induced with fibronectin or laminin (29). In the lung cancer cell line A549, EMT is induced via TGF-β stimulation (30, 31) or adhesion to Col-I (32). Given the protocol described (32), the Col-I substrate used appeared to be a non-gel-form, which piqued our curiosity in studying the effect of Col-I gels on EMT induction.

Cadherins are the transmembrane components of the adherence junction at cell–cell contact sites, comprising an extracellular domain responsible for cell–cell homophilic interactions, a transmembrane domain, and a cytoplasmic tail that binds catenin (33). Catenin links cadherin to the actin cytoskeleton and functions in cellular signaling (33, 34). Cadherin molecules have multiple isoforms. E-cadherin is found in polarized epithelial cell–cell junctions widely, whereas E- and P-cadherin are found in squamous epithelial cell junctions. Notably, fibroblasts form intercellular junctions containing N-cadherin (33). The switching of cadherin from the epithelial to the mesenchymal type (i.e., cadherin switching) is an important process in EMT induction (11–13, 35, 36). Many epithelial cells begin to express mesenchymal markers during EMT process, including vimentin (13, 37, 38) and fibronectin (37, 38). Cadherin switching and increased integrins are considered associated with cancer cell migration (13).

It is often assumed that cancer cells are anchorage-independent. Cancer cells are insensitive growth regulatory signals originating from adhesion to the ECM (39). However, accumulating evidence suggests that integrin-mediated contact of malignant cells to the ECM influences their behavior via signal transduction (40–42). Henriet *et al*. reported that human melanoma cells (M24met) cultured on collagen gels were growth-arrested at the G1/S checkpoint and maintained high levels of p27KIP1 mRNA and protein through β1 integrin-mediated mechanisms (42). We previously reported that adhesion to Col-I gels suppressed cell proliferation and PI3K/Akt pathway activation in three cancer cell lines (43). Akt activation is characteristically suppressed on collagen gels in these cells and keratinocytes (9,, 43, 44).

This study investigates the EMT-related changes in A549 cells cultured on gels and non-gel forms of Col-I and on non-treated culture dishes with and without TGF-β1. We assessed cell morphology, proliferation, single-cell migration and EMT-related features.

## Results

### Substrate-dependent cell morphology, cluster formation and proliferation of A549 cells

A549 cells were cultured in four culture conditions: on non-treated culture dish (on NT), on non-gel form of Col-I coated culture dish (on Col-I-non-gel), on fibrous Col-I gels coated culture dish (on Col-I-gel) and on non-treated culture dish with TGF-β1 in the culture media (with TGFβ1). The cells were seeded at a single-cell density (1.0 × 10^5^ cells / well of six-well culture plate) and cultured for 2 days. Cellular morphology was observed using phase-contrast microscopy. On NT, the cells showed typical epithelial cell clusters. On Col-I-non-gel, cell–cell interaction was weakened and the cell morphology was converted from an epithelial phenotype to a fibroblastic phenotype. The cell morphology on Col-I-gel differed from the previously mentioned two conditions. Cell spreading was suppressed, and some cells formed clusters (Fig.1A, arrows). The other cells did not form cell–cell contacts and maintained single-cell form (Fig.1A, arrow heads). With TGFβ1, A549 cells spread very well, showing a fibroblastic-like morphology and not forming cell clusters (Fig. 1A).

**Figure 1.**
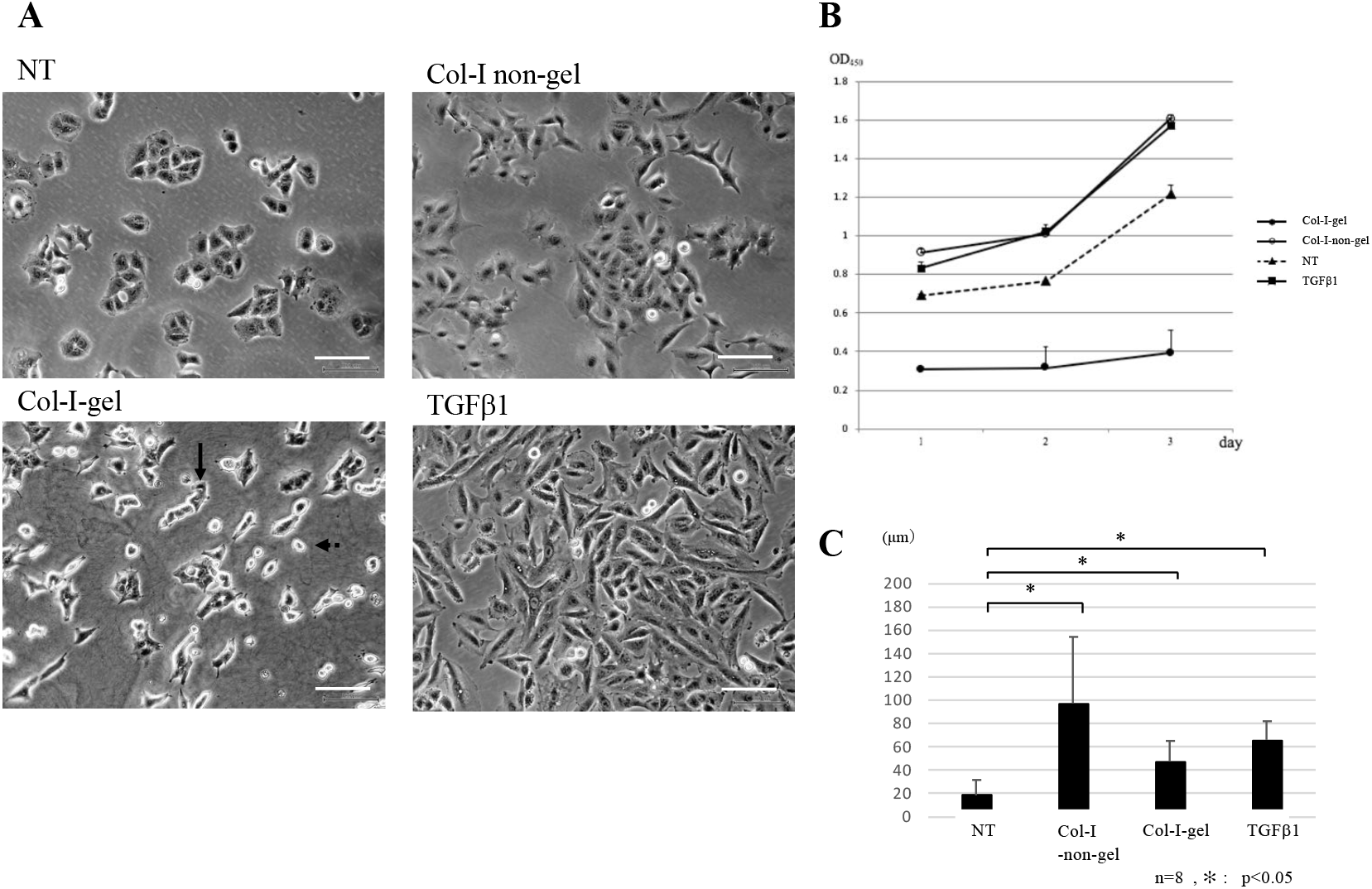
Morphology, proliferation and migration of A549 cells. **A**. A549 cells were cultured on NT, on Col-I-non-gel, on Col-I-gel and with TGFβ1 for 2 days. Cell morphology was observed using phase-contrast microscopy. Scale bars :100 µm **B**. The number of viable cells was estimated using the WST-8 method on days 1, 2, and 3. A549 cells were cultured on NT (dotted line, triangles), Col-I-non-gel (solid line, open circles), Col-I-gel (solid line, closed circles), and TGFβ1 (dotted line, crosses). The experiments were performed in triplicate, and data are shown as means ± SD. **C**. Monitoring of the migration of single cells for 2 days by using time-lapse microscopy. The relative distance of eight randomly selected cells on each condition was assessed. The mean relative distances are shown as means ± SD. Differences between two individual groups were analyzed using Student’s t-test. \**P*<0.05

Cell proliferation was estimated at the day 3 of culture. The cells continued proliferating on NT, on Col-I-non-gel, and with TGFβ1. Although cell proliferation on Col-I-gel was maintained, it was markedly suppressed compared with the other conditions (Fig. 1B).

### Effects of Col-I gel on the migration of A549 cells

We monitored A549 cells in previously mentioned four culture conditions using a time-lapse observation system to examine effects of Col-I substrates on single-cell migration. The relative distance (distance between tracking start position and current position) of eight randomly selected cells in each condition was analyzed to quantitatively estimate cell motility. A549 cells on NT exhibited limited migration. Cells hardly moved from the place where they were, proliferated and formed clusters. The mean relative distance was notably the shortest among the tested conditions (Fig. 1C, Supporting Information S-1). The cells on Col-I-non-gel spread, moved well, and did not form cell clusters. The mean relative distance was the longest among the tested conditions (Fig. 1C, Supporting Information S-2). A significant difference in the relative distance between the Col-I-non-gel and the NT groups was observed. Considering the cell image on Col-I-gel at 2 days-culture, we initially hypothesized that the cells would hardly migrate and proliferate to form clusters. Contrary to expectation, on Col-I-gel, some single-cells subsequently migrated to form cell clusters and the others did not move from the place where they were. In the cells on Col-I-gel (Fig. 1C, Supporting Information S-3) and with TGFβ1 (Fig. 1C, Supporting Information S-4), the mean relative distances fell between the above two conditions and those were significantly longer than that on NT. The mean relative distance with TGFβ1 was longer than that on Col-I-gel. Furthermore, there was a significant difference between the Col-I-gel and the NT groups and between the TGFβ1 and the NT groups.

### Switching of E- to N-cadherin was observed in A549 cells cultured on Col-I-gel as well as with TGFβ1

E-cadherin is the transmembrane components at cell–cell contact sites in epithelial cells (33). A549 cells cultured for 2 days were immunostained with anti-E- and N-cadherin antibodies. The cells on NT expressed E-cadherin but not N-cadherin in cell– cell contact regions. On Col-I-non-gel, both E- and N-cadherin were hardly observed in cell–cell contact regions. On Col-I-gel, E-cadherin was hardly observed, while, N-cadherin was observed in cell–cell contact regions. Single-cells expressed both cadherins like a dot. Following with TGFβ1, the cells spread well and showed slight cell–cell contact. These regions of slight contacts did not express E-cadherin, whereas N-cadherin was occasionally observed (Fig. 2A).

**Figure 2.**
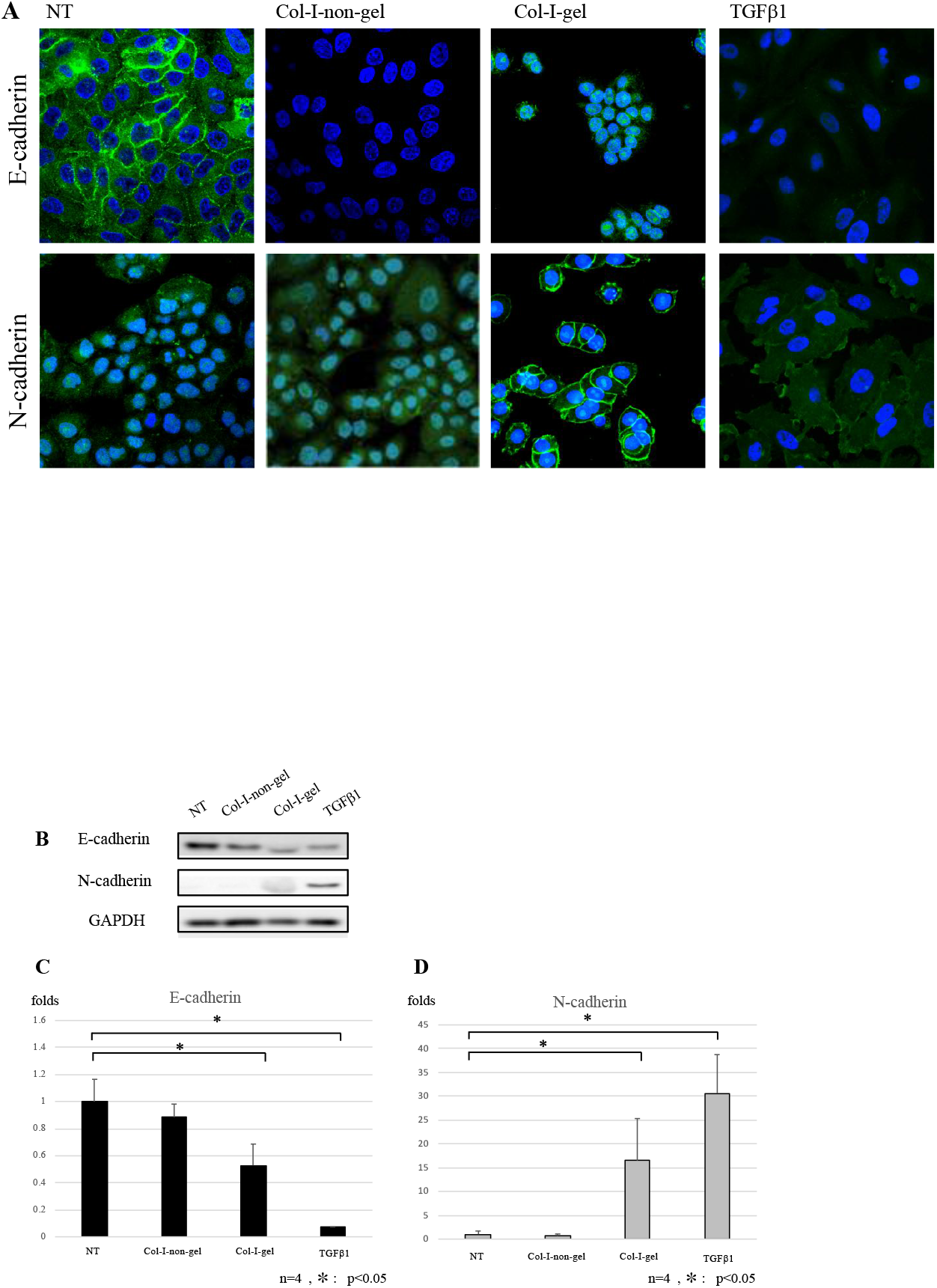
Expression of E-cadherin and N-cadherin in A549 cells. **A**. A549 cells were cultured on NT, on Col-I-non-gel, on Col-I-gel and with TGFβ1 for 2 days. Cells were stained with E- and N-cadherin antibodies and observed using a laser scanning microscope. **B**. Typical western blot data are shown. **C**. E-cadherin protein expression levels were determined from the western blots. **D**. N-cadherin protein expression levels were determined from the western blots. The mean values in each condition were divided by that of NT. Differences between two individual groups were analyzed using Student’s t-test. \**P*<0.05

Western blotting was performed to quantify the cadherin expression. Typical western blot data are presented in Figure 2B. Cadherin expression values were normalized to that of glyceraldehyde 3-phosphate dehydrogenase (GAPDH). The relative expression levels of each condition were estimated based on the expression on NT. The mean value of E-cadherin expression was the highest in cells on NT, followed by on Col-I-non-gel, on Col-I-gel, and with TGFβ1 (Fig. 2C). In contrast, the mean of N-cadherin expression was the lowest in cells on NT and increased in the order of on Col-I-non-gel, on Col-I-gel, and with TGFβ1 (Fig. 2D). E-cadherin expression levels on Col-I-gel and with TGFβ1 were significantly lower than that on NT. In the case of N-cadherin, those on Col-I-gel and with TGFβ1 were significantly higher than that on NT. Significant differences were observed in both E- and N-cadherin between the Col-I-gel and the NT groups and between the TGFβ1 and on NT groups (Fig.2C, 2D). The western blotting results did not coincide with the immunohistochemical observations of E-cadherin on Col-I-non-gel or N-cadherin with TGFβ1. We hypothesized this was caused by loose cell–cell contacts due to scattered cells (Fig.1A). Thus, cadherins might not accumulate and would be invisible by immunohistochemical analysis. A considerable amount of E-cadherin expression was observed by western blotting on Col-I-gel, but was not observed by immunostaining. The reason for this is unclear, it could be because of the mixture of single cells and cell clusters on Col-I-gel (Fig. 1A).

### Vimentin and integrins did not increase in cells on Col-I-gel, unlike those with TGFβ1

Vimentin is a type III intermediate filament found in mesenchymal cells, maintaining cell and tissue integrity (45). Increasing vimentin expression enhances directed cell migration, and the vimentin network regulates the actin cytoskeleton, which is associated with cell migration (46). We stained cells cultured for 2 days in each condition with anti-vimentin antibody. Immunostaining did not show vimentin in A549 cells on NT, on Col-I-non-gel and on Col-I-gel. In contrast, vimentin was present in the spreading A549 cells with TGFβ1 (Fig. 3A). Typical western blot data are presented in Figure 3B. Vimentin expression was measured and normalized to that of GAPDH. The expression levels of each condition were estimated based on that on NT. Western blottig analysis corroborated the immunohistochemical observations. A significant difference between the Col-I-gel and the TGFβ1 groups and between the TGFβ1 and the NT groups was observed (Fig. 3C).

**Figure 3.**
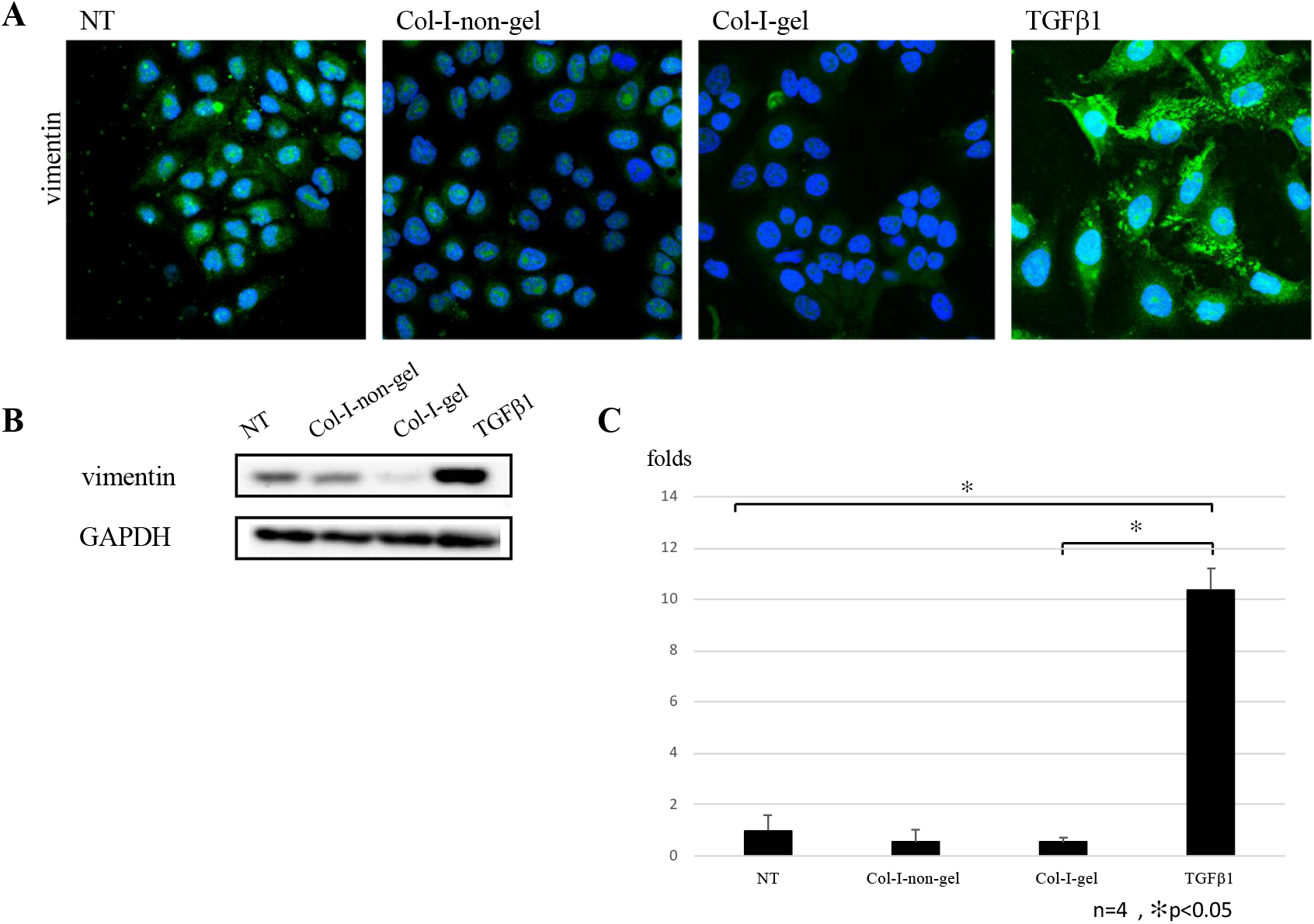
Expression of vimentin in A549 cells. **A**. A549 cells were cultured on NT, on Col-I-non-gel, on Col-I-gel and with TGFβ1 for 2 days. Cells were immunostained with anti-vimentin antibody. **B**. Typical western blot data are shown. **C**. The levels of protein expression were determined from the western blots and normalized to that of GAPDH. The mean values of vimentin in each condition were divided by that on NT. The results were verified by Student’s *t*-test. \**P* < 0.05.

Integrins are heterodimeric cell surface receptors that composed of α and β subunits. They bind between the ECM and the inside of the cytoskeleton (47, 48). Integrin α1β1, α2β1, α10β1, and α11β1 are reported as collagen receptors. Most integrins have intracellular linkages to the actin cytoskeleton (47). Integrins are thought to contribute to the cancer cell properties of limitless proliferation, invasion, promotion of tumor angiogenesis, evasion of apoptosis, and the development of resistance to growth suppressors (48). Immunostaining of the cell–cell contact regions and in cell peripheral regions of A549 cellscultured on NT, on Col-I-non-gel, and on Col-I-gel showed integrin β1. Actin was also observed in the peripheral regions of cells. The stained images of integrin β1 and actin on NT, on Col-I-non-gel, and on Col-I-gel were similar. With TGFβ1, the cells spread well, and notably, actin filaments developed as long straight lines (arrows, Fig. 4A). The cellular expression of integrin β1 and α2 was measured using flow cytometry. The expression ratio of integrins was standardized against the mean values of the fluorescein intensity of cells cultured on NT. In both cells cultured on Col-I-non-gel and on Col-I-gel, the mean integrin β1 and α2 expression values were the same as those on NT (Fig. 4B and Table 1). By contrast, the mean values of integrins β1 and α2 increased by 1.13-fold and 1.94-fold, respectively, with TGFβ1 (Fig. 4C and Table 1).

**Table 1.**
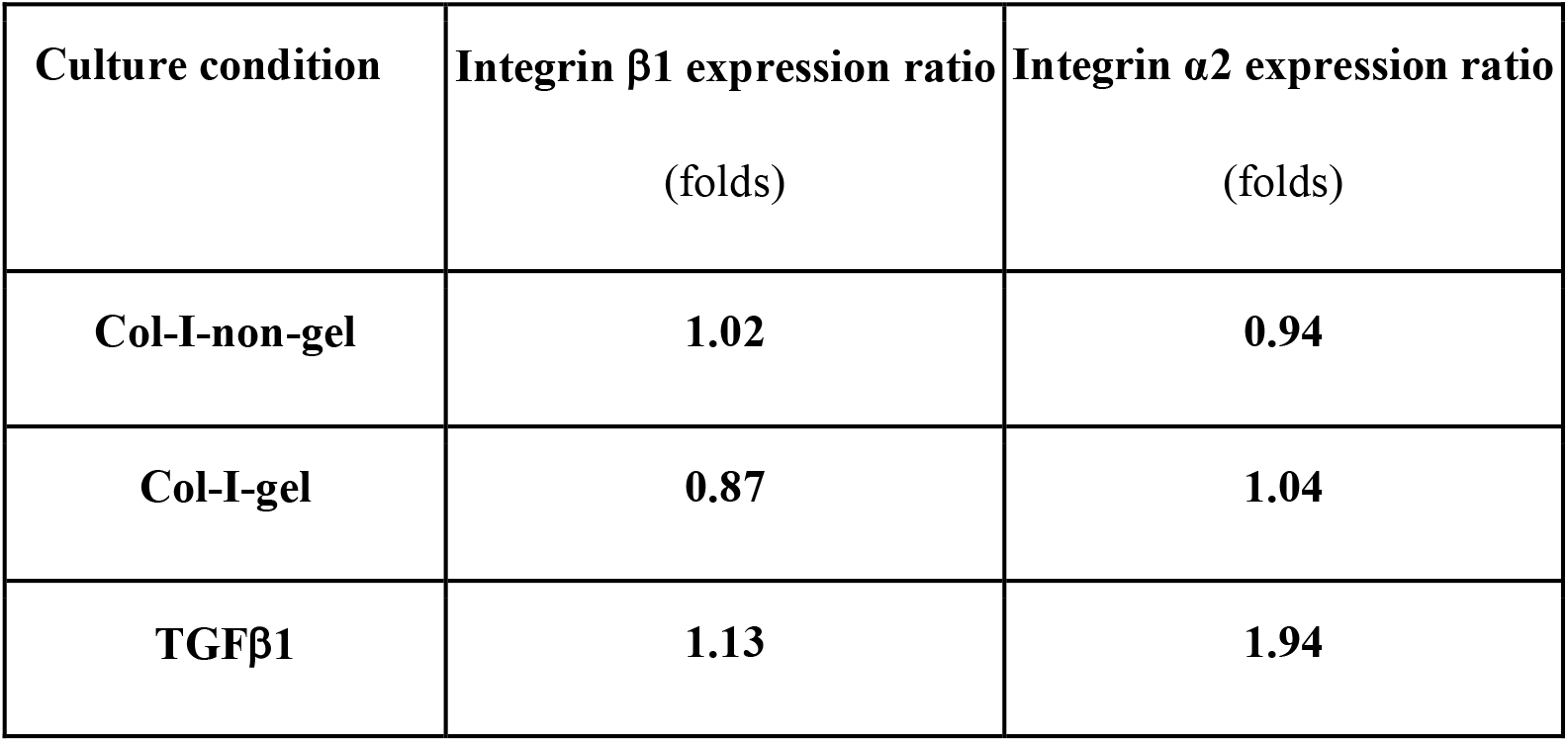
Comparative integrin expression ratio in each culture condition comparing with NT

| **Culture condition** | **Integrin β1 expression ratio**<br>(folds) | **Integrin α2 expression ratio**<br>(folds) |
| --- | --- | --- |
| **Col-I-non-gel** | **1.02** | **0.94** |
| **Col-I-gel** | **0.87** | **1.04** |
| **TGFβ1** | **1.13** | **1.94** |

**Figure 4.**
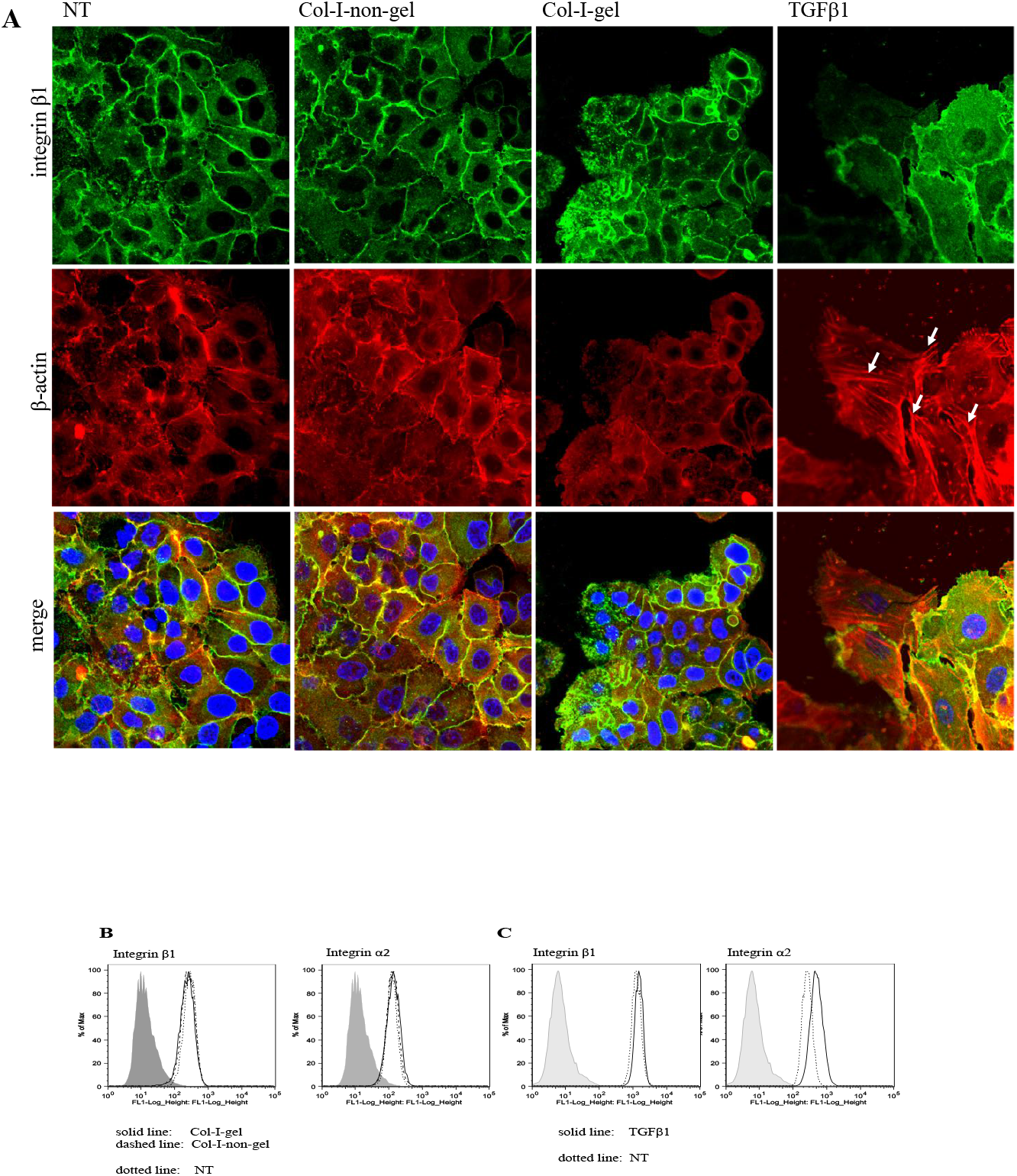
Expression of integrins β1 and α2 in A549 cells. **A**. Integrin β1 and β-actin were observed by immunostaining. **B**. The fluorescence intensity of integrins β1 and α2 were estimated by flow cytometric analysis. Comparing with the data on NT, on Col-I-non-gel, and on Col-I-gel. Gray-filled line, unstained cells (negative control); dotted line, on NT; dashed line, on Col-I-non-gel; solid line, on Col-I-gel. **C**. Comparing with the data on NT and with TGFβ1. Gray-filled line, unstained cells (negative control); dotted line, NT; solid line, with TGFβ1.

## Discussion

In this study, we demonstrated that adhesion to Col-I fibrous gels induced a unique transition in A549 cells, i.e. E- to N-cadherin switching, forming cell clusters via N-cadherin and upregulation of single-cell motility as compared with NT. The mean value of relative distance on Col-I-gel was shorter than that with TGFβ1 or on Col-I-non-gel. Such behaviors differed from that of with TGFβ1, as a typical EMT-induced condition. The similarities and differences of A549 cells between on Col-I-gel and with TGFβ1 are summarized in Table 2.

**Table 2.**
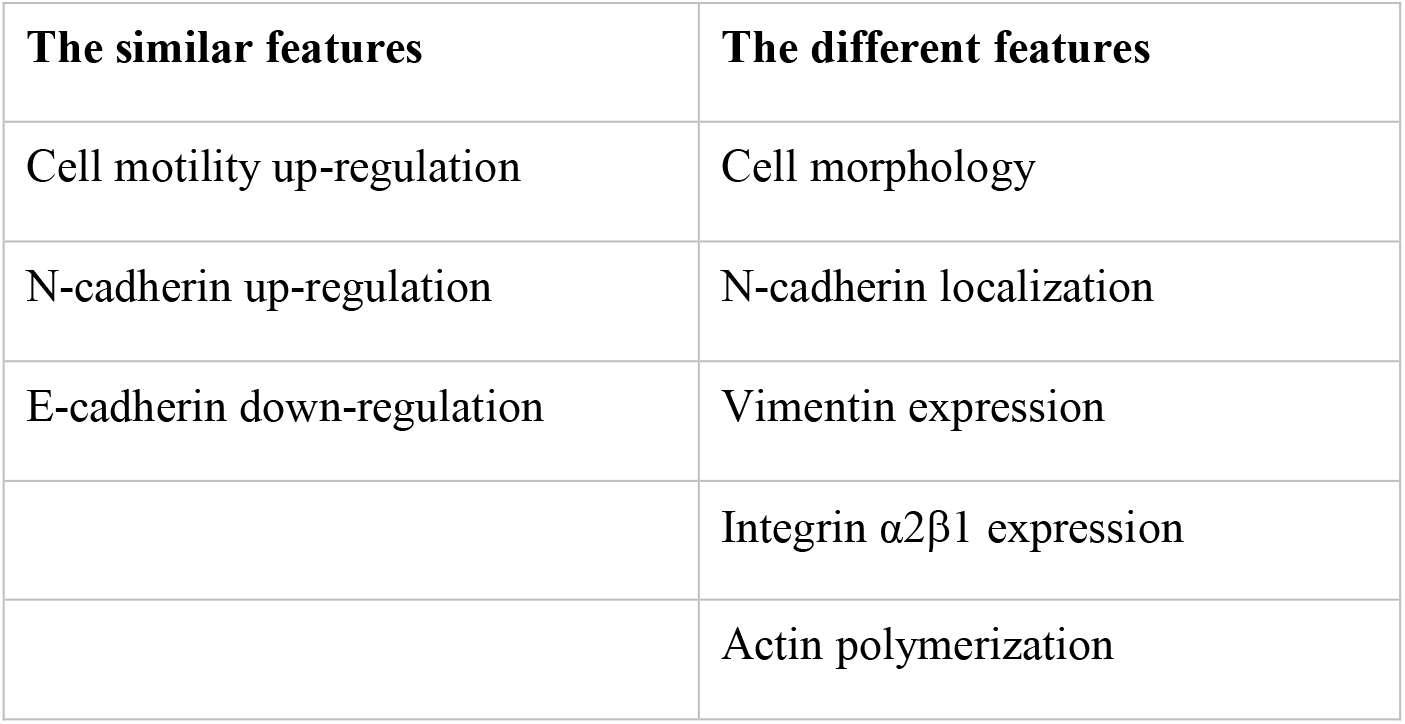
Comparative cell features between on Col-I-gel and with TGFβ1 The features of A549 cells cultured on Col-I-gel and with TGFβ1 are summarized. Culture conditions: A549 cells cultured for 2 days on Col-I gels (Col-I-gel), on non-treated culture dish with TGF-β1 (with TGFβ1).

| The similar features | The different features |
| --- | --- |
| Cell motility up-regulation | Cell morphology |
| N-cadherin up-regulation | N-cadherin localization |
| E-cadherin down-regulation | Vimentin expression |
| | Integrin α2β1 expression |
|  | Actin polymerization |

A few papers have been reported for EMT induction by adhesion to Col-I in A549 cells (32, 49). Shintani et al. showed that adhesion to Col-I promoted single-cell scattering. Their results were close to that of our Col-I-non-gel condition, but there were some differences (32). We think that the main cause of this is due to the different Col-I treatment protocols. Since Shintani et al. did not mention the Col-I treatment protocols in detail, the gel-forming states were unknown (29, 32, 35). Because Col-I forms fibrils under physiological conditions, Col-I molecules coated on culture dishes may form fibrils during the cell culture period. Even if gels may not form, it is highly possible to form fibrils depends on the amount of applied Col-I. Thus, a different Col-I treatment protocol would lead to different fibril-forming states on the culture dish. However, Col-I treatment protocols are not unified in general, and the state of fibril formation on culture dishes has received little attention.

Gel culture systems utilizing Col-I fibrils are thought to be better to mimic the physiological environment than those using the non-gel form of Col-I (50, 51). Considering the published papers, there are at least three essential difference between fibrous Col-I gels and Col-I-non-gel form as culture substrates (52–54). First, the specific integrin-recognition sites on collagen molecules are masked on fibrils (52). Second, the matrix stiffness of gel-formed Col-I fibrils is lower than that of the plastic dish surface (53). Third, the morphology of Col-I gel surface is porous and not flat (54). Baker and coworkers have reported differences in cell morphology and proliferation on three-dimensional gel culture systems between flat hydrogel (e.g., polyacrylamide gels) and synthetic fibrous materials (e.g., using electrospinning) (54). Each experimental system in these papers (52–54) is designed to focus on a single factor, but these factors can collectively affect cells in Col-I gel culture system. We think it is crucial to study the phenomena in Col-I gel culture system to elucidate comprehensive roles of Col-I in the living body. For example, increasing matrix stiffness induces EMT (55, 56). In contrast, we observed that the switching from E- to N-cadherin was more prominent on softer Col-I-gel than that on harder Col-I-non-gel (Fig. 2), suggesting that the effect of matrix stiffness is important but not an only determinant factor to induce cadherin switching. Shukla et al. reported EMT induction and cell migration in A549 cells focusing on the effects of Col-I substrate stiffness. They examined cell migration parameters and cadherin switching on collagen-coated polydimethylsiloxane gels with varying degrees of stiffness. The migration parameters depended on substrate stiffness, while E- to N-cadherin switching was unaffected (49).

A correlation between N-cadherin expression and cell motility is now under debate. N-cadherin expression is upregulated in some invasive cancer cell lines (57, 58). N-cadherin overexpression induces an invasive morphology in squamous tumor cells (59) and breast cancer cells (60) *in vitro*. N-cadherin knockdown cells exhibit reduced migration and a limited ability to scatter on collagen in pancreatic cancer cells, and N-cadherin expression promote metastasis in an orthotopic model (29). However, there is no correlation between N-cadherin expression and invasion in astrocytoma and glioblastoma (61, 62). In our study, there was no simple correlation between N-cadherin expression (Fig. 2D) and cell migration (Fig. 1C). We think this is caused by the balance of the following two factors. Signal activation via adhesion to cell recognition sites on Col-I fibrils promotes cadherin switching induction and migration in cells. On the other hand, gel natures of Col-I, e. g., gel stiffness and an uneven surface condition might suppress cell migration. Considering each factor, more study is necessary to elucidate the correlation between cadherin switching and cell motility on fibrous materials like Col-I gels.

In EMT induction by TGF-β signaling activation, EMT-related signals often lead and enhance TGF-β autocrine. TGF-β autocrine sometimes goes into a closed-loop system (autocrine loop), and it contributes to enhance and maintain the mesenchymal state in cells (25, 63, 64). Once TGF-β signaling activates, TGF-β autocrine activation maintains via TGF-β/ZEB/miR-200 signaling network to keep the mesenchymal phenotype in MDCK cells (63). Sphingosine-1-phosphate-induced EMT is mediated by syndecan-1 activation and TGF-β autocrine (64). In our study, the expression of mesenchymal cytoskeleton (vimentin) and collagen receptor (integrin α2β1) was increased in TGF-β1-treated cells. In contrast, they did not increase on both states of Col-I (Fig. 3, 4). Thus, we considered that TGF-β signal activation did not affect cells on Col-I. The development of the cytoskeleton and integrin– ECM interaction enhances cell migration (65).

The interstitial Col-I fibril lattices function as both a cell scaffold and a barrier to prevent cell migration in cancer invasion process (6, 66). Furthermore, EMT induction via adhesion to Col-I is thought to enhance cancer invasion (32). The effects of adhesion to Col-I (regardless of the state of fibril formation) are still ambiguous. N-cadherin-mediated cell cluster formation in cadherin switching induced-cells is a unique phenomenon. It may provide the key to understand as yet unelucidated biological and pathological phenomena. A schematic view of the roles of Col-I fibrils in cancer invasion is shown in Figure 5.

**Figure 5.**
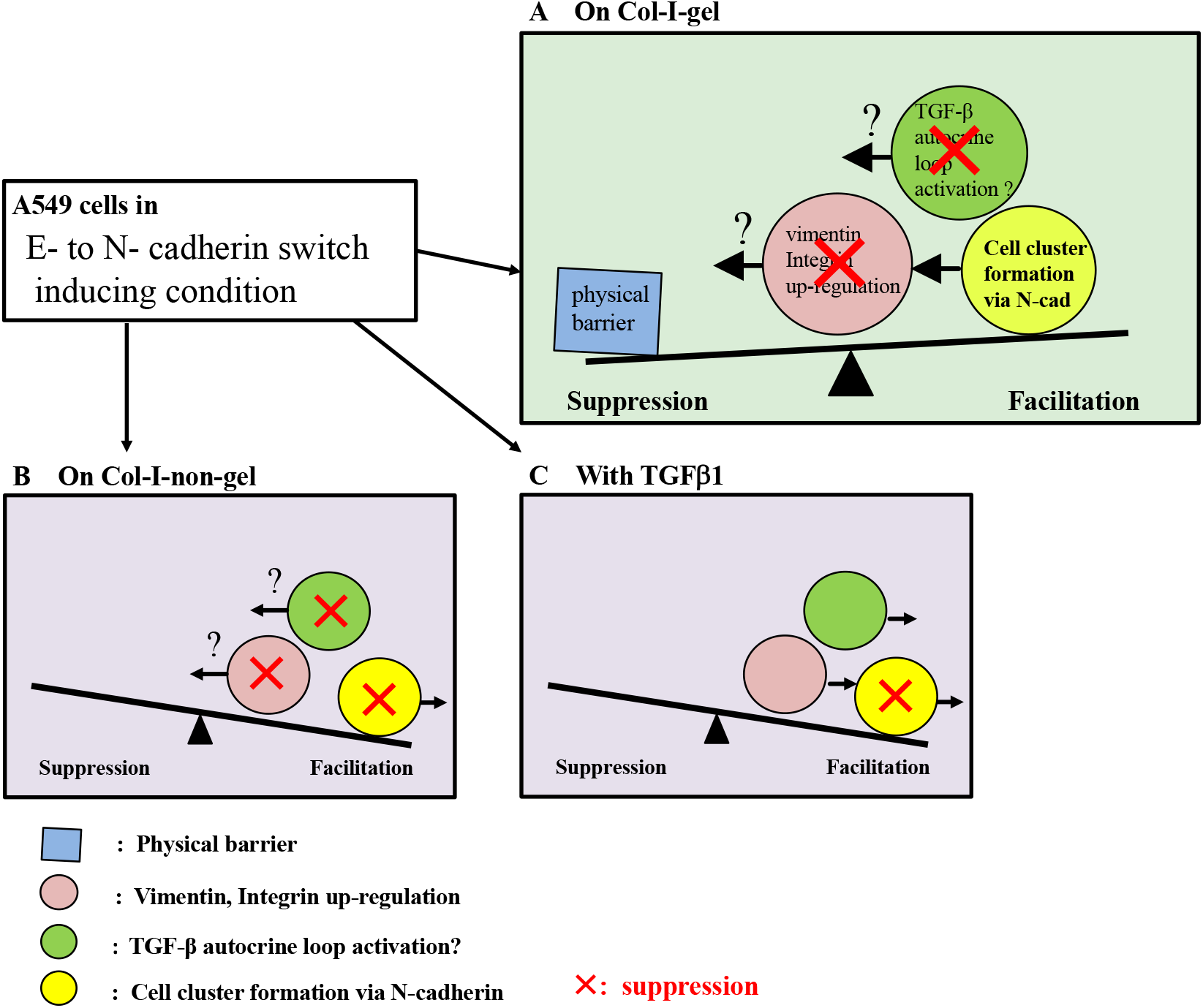
Schematic view of the roles of Col-I fibrous gels on cancer invasion process. **A**. On Col-I-gel, E- to N-cadherin switching is induced and cell clusters are formed via N-cadherin in A549 cells. The expression levels of mesenchymal intermediate filament (vimentin) and collagen receptor (integrin α2β1) do not increase comparing with on NT. Thus, TGF-β autocrine loop might not work. In addition, Col-I fibril lattices might work as a barrier to prevent cell invasion. These changes might work as candidates for suppressor in cancer cell invasion. **B**. On Col-I-non-gel, unchanged of the expression levels of vimentin and integrin α2β1, and TGF-β autocrine loop suppression work as invasion suppressors. On the other hand, there are neither cell cluster formation via N-cadherin nor barrier of Col-I fibril lattice. **C**. With TGFβ1, all of these factors on Col-I-gel do not work.

Many individuals may have undiagnosed tumors in their bodies for several years (67, 68). Bissell and Hines reported that the microenvironment surrounding the tumors was the source of tumor-suppressive signals in these cases (68). Thus, Col-I fibrils may be one of the niche factors affecting cancer dormancy.

## Experimental procedures

### Cell culture

Human lung cancer cell line A549 cells were provided by the RIKEN BRC through the National Bio-Resource Project of the Next Japan. The cells were maintained in Dulbecco’s modified Eagle Medium (DMEM; Sigma-Aldrich, St. Louis, MO, USA) supplemented with 10% fetal bovine serum (FBS; Biowest, Nuaille, France).

### TGF-β1 treatment

A549 cells were treated with 2 ng/ml TGF-β1 (Sigma-Aldrich, Merck KGaA, Darmstadt, Germany) for 2 days in DMEM with 10% FBS on non-treated culture dish surfaces (31) to induce EMT.

### Preparation of culture substrates

Acid-soluble bovine Col-I was obtained from Nippi Inc. (Tokyo, Japan). The plastic surfaces of tissue culture plates were treated with Col-I in two ways as previously described (9). For the non-gel form, the wells were coated with Col-I (10 μg/ml) in 1 mM HCl for 1 h at room temperature. For the gel form, 1 mg/ml Col-I solution in PBS (–) (phosphate-buffered saline without calcium and magnesium) was added to the wells of a culture plate (96-well plate, 0.1 ml/well; six-well plate, 1 ml/well) and incubated for 1 h at 37 °C in a humidified CO2 incubator (9, 43).

### Analysis of cell proliferation

The number of living cells was estimated using a WST-8 (modified tetrazolium salt) cell proliferation kit (Cell Counting Kit-8; Dojin, Kumamoto, Japan) according to the manufacturer’s protocol. Briefly, A549 cells (5.0 × 10^3^ cells/well) were cultured using DMEM including 10% FBS in a 96-well tissue culture plate treated with or without Col-I. The absorbance was measured at 450 nm using SH-9000 Lab microplate reader (Corona Electric Co., Ltd., Ibaraki, Japan). Each condition was assessed in triplicate and the results were expressed as means ± standard deviation (SD).

### Analysis of cell migration using time-lapse microscopy

The migration of A549 cells was monitored using the time-lapse function of BZ-X800 microscope system (Keyence, Osaka, Japan) housed in a 37 °C temperature-controlled chamber. Culture plates (six-well) were used in the non-treated condition or preincubated with Col-I (gels or non-gel-form). The cells were seeded at a concentration of 1.0 × 10^5^ cells/well on the surface of both forms of Col-I and on non-treated dish surfaces with or without TGF-β1 (2 ng/ml) and preincubated in a CO2 incubator for 2 h. Subsequently, the cells were placed in a microscope chamber and monitored for 48 h. The relative distance (distance between the tracking start position and the current position) of eight randomly selected cells in each condition was assessed using the BZ-H4K Motion Analysis Application (Keyence, Japan), expressed as means ± SD. Calculations were performed using Microsoft Excel for Mac (ver. 16.16.24). Differences between two individual groups were analyzed using Student’s t-test (two-tailed). Results were considered significantly different when *P* < 0.05.

### Antibodies

Mouse monoclonal E-cadherin antibody (HECD-1) was purchased from Takara Bio Inc. (Shiga, Japan). Rabbit monoclonal N-cadherin (D4R1H), vimentin (D21H3), and GAPDH (14C10) antibodies were purchased from Cell Signaling Technology, Inc. (Beverly, MA, USA). Mouse monoclonal integrin β1-activating (TS2 /16) and mouse monoclonal integrin α2 (P4B4) antibodies were purchased from Santa Cruz Biotechnology, Inc. (Dallas, TX, USA). Peroxidase-conjugated secondary antibodies against rabbit IgG and mouse IgG were purchased from Cell Signaling Technology, Inc. The Alexa Fluor 488-conjugated antibodies against mouse IgG and rabbit IgG were obtained from Santa Cruz Biotechnology, Inc. Phalloidin-tetramethylrhodamine B isothiocyanate was obtained from Sigma-Aldrich.

### Immunohistochemical analysis

Prior to cell culture, twelve-well Teflon-coated slides (Thermo Fisher Scientific, Waltham, MA, USA) were treated with Col-I as previously described (9). Following the growth of cells on the slides for 2 days on each condition, they were washed with PBS (–) and fixed with 4% paraformaldehyde for 1 h at room temperature. Then, the cells were washed with PBS (–), permeabilized, incubated in 1% FBS/PBS (–) for 1 h at room temperature, and incubated with monoclonal antibodies overnight at 4 °C. The cells were washed thrice with PBS-Tween 20 and incubated with Alexa Fluor 488-conjugated secondary antibody (diluted 1: 200 in PBS (–)) for 1 h. Finally, the cells were stained with VECTASHIELD Antifade Mounting Medium with DAPI (Vector Laboratories, Burlingame, CA, USA) and observed using an FV1200 laser scanning microscope (Olympus, Tokyo, Japan).

### Western blotting

Prior to the preparation of samples for western blot analysis, A549 cells (1.0 × 10^5^ cells/well) in six-well tissue culture plates containing 1 ml DMEM were cultured for 2 days on each condition. In the case of Col-I-gel, the cells and gels were gathered together and centrifuged. After discarding the supernatant, the pellet was washed with PBS (–) and incubated on ice for 5 min in 0.1 ml cell lysis buffer (Cell Signaling Technology, Inc.). In the other conditions, the cells were washed with PBS (–) and lysed on ice for 5 min in 0.05 ml cell lysis buffer as previously described (9). The samples were electrophoresed on a 10% polyacrylamide gel and electrophoretically transferred to nitrocellulose membranes. The membranes were blocked overnight at 4 °C with 2% nonfat dry milk in tris-buffered saline (TBS) containing 0.05% Tween-20 (TBS-T), followed by incubation with primary antibody diluted in TBS-T. The bands were visualized using Amersham ECL Prime Western Blotting Detection Reagents (GE Healthcare, Amersham, UK) according to the manufacturer’s instructions. Four independent experiments were conducted. The protein concentrations were determined using Fiji software (Media Cybernetics, Rockville, MD), and expressed as means ± SD. Calculations were performed using Microsoft Excel for Mac (ver. 16.16.24). Differences between two individual groups were analyzed using Student’s t-test (two-tailed). Results were considered significantly different when *P* < 0.05.

### Flow cytometry

Prior to the flow cytometric analysis, A549 cells (1.0 × 10^5^ cells/well) were seeded in six-well tissue culture plates and cultured on each condition for 2 days. On Col-I-gel, the cells were gathered from Col-I gels using 1 mg/ml collagenase (Wako, Japan) at 37 °C for 5 min and then centrifuged and discarded supernatant. Precipitated cells were separated into single cells by treatment with 0.25% trypsin-0.02% (trypsin-EDTA) (Novozymes North America, Inc., Franklinton, NC, USA) at 37 °C for 5 min. In the other conditions, the cells were treated with 1 mg/ml collagenase (Wako) at 37 °C for 5 min and gathered with trypsin-EDTA treatment at 37 °C for 5 min. Then the cells were resuspended in the treatment buffer and stained with anti-bodies as previously described (69). Immunofluorescent-stained cells were analyzed using a FACSAria II Flow Cytometer (BD Bioscience, Franklin Lakes, NJ, USA) and the resultant data were analyzed using the FlowJo software (Tomy Digital Biology Co., Ltd., Tokyo, Japan). More than 5,000 cells were examined for each condition.

## Data availability

The datasets available from the corresponding author on request.

## Acknowledgement

This work was performed in part under the Cooperative Research Program of Institute for Protein Research, Osaka University (CR-15-01-7).

The authors would like to thank Enago (www.enago.jp) for the English language review.

## Footnotes

## The abbreviations

Col-I: type I collagen
Col-I-gel: fibrous Col-I gels
Col-I-non-gel: non-gel form of Col-I
DMEM: Dulbecco’s Modified Eagle Medium
DMEM (10): DMEM supplemented with 10% fetal bovine serum
ECM: extracellular matrix
EMP: Epithelial-mesenchymal plasticity
EMT: Epithelial-to-mesenchymal transition
FBS: fetal bovine serum
GAPDH: glyceraldehyde 3-phosphate dehydrogenase
NT: non-treated cell-culture dish
PBS (-): phosphate-buffered saline without calcium and magnesium
PBS-T: PBS (-) containing 0.05 % Tween 20
PI3K: phosphatidylinositol 3-kinase
ROS: reactive oxygen species
TBS: tris-buffered saline
TBS-T: TBS containing 0.05% Tween-20
TGF-β: transforming growth factor-β
TGFβ1: non-treated culture dish with TGF-β1 in the culture media

